# Identification a novel subset of alveolar type 2 cells expanding following pneumonectomy and enriched in PD-L1

**DOI:** 10.1101/2020.05.20.106005

**Authors:** Negah Ahmadvand, Farhad Khosravi, Arun Lingampally, Roxana Wasnick, Ivonne Vazquez-Armendariz, Monika Heiner, Stefano Rivetti, Yuqing Lv, Jochen Wilhelm, Andreas Gunther, Susanne Herold, Denise Al Alam, Chengshui Chen, Parviz Minoo, Jin-San Zhang, Saverio Bellusci

## Abstract

Alveolar type 2 (AT2) cells are heterogeneous cells; where specialized AT2 subpopulations within this lineage exhibit stem cell properties. However, the existence of quiescent, immature cells within the AT2 lineage, which get activated during lung regeneration, is unknown.

*Sftpc^CreERT2/+^; tdTomato^flox/flox^* mice were used for the labelling of AT2 cells and labeled subpopulations were analyzed by flow cytometry, qPCR, ATAC-seq, gene arrays, pneumonectomy, and culture of precision-cut lung slides. Human lungs from donor and IPF were also analyzed.

In mice, we detected two distinct AT2 subpopulations with low tdTomato level (Tom^Low^) and high tdTomato level (Tom^High^). Tom^Low^ express lower level of AT2 differentiation markers, *Fgfr2b* and *Etv5*, while Tom^High^, as *bona fide* mature AT2 cells, show higher level of *Sftpc*, *Sftpb*, *Sftpa1*, *Fgfr2b*, and *Etv5*. ATAC-seq analysis indicates that Tom^Low^ and Tom^High^ constitute two distinct cell populations with specific silencing of *Sftpc*, *Rosa26* and cell cycle gene loci in Tom^Low^. Upon pneumonectomy, Tom^Low^ but not Tom^High^ cells proliferate and upregulate the expression of *Fgfr2b*, *Etv5*, *Sftpc*, *Ccnd1* and *Ccnd2* compared to sham. Tom^Low^ cells overexpress PD-L1, an immune inhibitory membrane receptor ligand, which is used by flow cytometry to differentially isolate these two sub-populations. In the human lung, PD-L1 and HTII-280 antibodies are used by flow cytometry to differentially sort mature AT2 (HTII-280-high, PD-L1-low) as well as an additional subpopulation of epithelial cells characterized by HTII-280-Low and PD-L1-high.

We have identified a novel population of AT2 quiescent immature progenitor cells in mouse that proliferate upon pneumonectomy and provided evidence for the existence of such cells in human.

**Significance of the work:** The characterization and mechanism of the proliferation of a novel and relevant pool of AT2 progenitor cells for the repair/regeneration process after injury are critical to improving respiratory function in patients with lung disease.

## INTRODUCTION

Alveolar type 2 (AT2) progenitor cells display self-renewal capabilities to maintain the AT2 cell pool and can also differentiate into AT1 cells following lung injury [1][2][3][4][5]. However, it is still debatable whether quiescent and immature AT2 progenitor cells exist in mouse and human lungs and if surface markers can be used to isolate these cells.

In rodents, AT2 cells are heterogeneous. Different subpopulations within this lineage, with different stem cell properties, have been identified based on various markers such as E-cadherin, Axin2, integrin σ6β4 and the dual expression of Surfactant protein C (Sftpc)/ Secretoglobin family 1A member a (Scgb1a1) [6][7][8][9]. AT2 cells can be divided equally into E-cadherin positive and negative cells. Upon hyperoxia injury, E-cadherin negative cells are more resistant, more proliferative and display higher level of telomerase activity, while E-cadherin positive cells are more sensitive to damage [6]. In the adult mouse lung, Axin2-positive cells make up 1% of the mature AT2 (Sftpc-high) cells and are distributed throughout the lung [8]. This ratio of Axin2-positive cells is stable over time. *In vivo* lineage tracing of Axin2-positive (Sftpc-high) cells revealed their ability to form clones of 2 to 5 cells accompanied by their expansion and differentiation into AT1 expressing Podoplanin (Pdpn). In addition, these cells display enhanced self-renewal capabilities in alveolosphere assays compared to the Axin2-negative AT2 cells suggesting that these Axin2-positive cells represent a subpopulation of mature AT2 cells with enhanced stem cell capacities [8]. In a similar and independent study, Axin2-positive cells belonging to the mature AT2 cell population were characterized. These cells, called alveolar epithelial progenitor (AEP), comprise around 20% of adult mature AT2 cells and are enriched for Wnt target genes and developmental genes such as *Fgfr2*, *Nkx2.1*, *Id2*, *Etv4*, *Etv5*, and *Foxa1.* These cells also display increased self-renewal ability in alveolospheres compared to the total AT2 cells [9].

Interestingly, AT2 cells heterogeneity is not limited to the adult lung. During the alveologenesis phase of lung development, Axin2-positive AT2 cells display enhanced proliferation compared to total AT2 cells [10]. A subset of Sftpc-negative, laminin receptor integrin α6β4-positive cells, located at the bronchoalveolar duct junctions (BADJ) and within the alveoli, regenerate AT2 Sftpc+ cells in the alveoli after severe lung injury [7]. Finally, bronchoalveolar stem cells, positive for the alveolar marker *Sftpc* and the club cell marker *Scgb1a1* are located at the BADJ and are amplified upon injury to give rise to either alveolar epithelial cells or club cells [11][12][13]. AT2 cells heterogeneity was equally shown in the context of diphtheria toxin (DT) injury as a subpopulation of lineage-labelled AT2 cells displaying relatively more resistance to DT toxicity was capable to repopulate the AT2 pool following injury. The identity of these survivor cells, however, is not clear [3].

In the present study, we identified a novel population of AT2 quiescent and immature progenitor cells characterized by low Sftpc level and high expression of the surface marker PD-L1, an immune checkpoint protein expressed in cancer stem cells and mediating anti-tumour suppression response [14][15][16]. We deployed a series of *in vivo* approaches in mice to label different subpopulations of AT2 cells, examined their epigenetic and transcriptomic characteristics and tested their respective response *in vivo* during lung regeneration following pneumonectomy as well as *in vitro* using the culture of precision-cut lung slices. We have also validated in mice and human the use of PD-L1 antibodies to differentially sort epithelial subpopulations. Our work opens the way for an in-depth characterization of this novel quiescent and immature AT2 stem cell population.

## MATERIALS AND METHODS

### Animal experiments

All animal studies were performed according to protocols approved by the Animal Ethics Committee of the Regierungspraesidium Giessen (permit numbers: G7/2017– No.844-GP and G11/2019–No. 931-GP).

### Human specimens

Human lung tissues from idiopathic pulmonary fibrosis (IPF) patients undergoing lung transplantation and non-IPF donors were provided from the Giessen biobank. The study protocol (AZ31/93) was approved by the ethics committee of University of Giessen.

### Lung dissociation and FACS

Single cell suspension was generated from adult lungs and stained with antibodies: anti-EpCAM (APC-Cy7-conjugated, Biolegend,1:50), CD49F (APC-conjugated, Biolegend,1:50), anti-PDPN (FITC-conjugated, Biolegend, 1:20), and anti-PD-L1 (unconjugated, Thermo Fisher, 1:100) antibodies for 20 minutes on ice in the dark, followed by washing. Then, the cells were stained for goat anti rabbit secondary antibody Alexa flour 488 (Invitrogen,1:500) for 20 minutes on ice in the dark. Next, cells were washed and stained for with SYTOX (Invitrogen) a live/dead cell stain according to the manufacturer’s instructions. Flow cytometry data acquisition and cell sorting were carried out using FACSAria III cell sorter (BD Biosciences, San Jose/CA). Data were analyzed using FlowJo software version X (FlowJo, LLC).

See online supplementary data for the human and mouse lungs dissociation.

### RNA extraction and quantitative real-time PCR

See online supplementary data

### Immunofluorescent Staining

See online supplementary data

### Alveolosphere assay

See online supplementary data

### Microarray

See online supplementary data

### ATAC-seq

See online supplementary data

### Quantification and statistical analysis

For quantification of immunofluorescence, cells were counted in 10 independent 63x fields per sample. Statistical analysis and graph assembly were carried out using GraphPad Prism 6 (GraphPad Prism Software). Significance was determined by unpaired two-tailed Student’s t-tests. Data are presented as mean ± standard error of mean (SEM). Values of *p* < 0.05 were considered significant. The number of biological samples (n) for each group are stated in the corresponding figure legends.

## RESULTS

### Identification of two AT2 subpopulations with different Sftpc and Fgfr2b levels

*Sftpc^CreERT2/+^; tdTomato^flox/flox^* mice were used to lineage label AT2 cells in the adult lung. In our experimental approach, tamoxifen was delivered in the water for one week. Two distinct populations of tdTomato^Pos^ cells were identified using flow cytometry analysis of tdTomato^Pos^ cells (Figure 1a). Of note, a similar observation, which was not followed up, was previously reported using the *Sftpc^Cre-ER^* mouse line generated by the Hogan lab [17]. In average, the tdTomato^Low^ (Tom^Low^) represented 9.93% of the overall Epcam^Pos^ cells (9.93% ± 1.73%, n=4) and the tdTomato^High^ (Tom^High^) represented 42.75% of the overall Epcam^Pos^ cells (42.75% ± 1.22%, n=4). We also confirmed that in this mouse line, epithelial cells were specifically labeled and that it mostly targeted AT2 Sftpc^Pos^ cells (Figure S1a,b). Very few BACS cells, identified through their localization at the BADJ, are labeled (0.1% of the total Tom^Pos^ cells), consistent with the literature [7]. Our labeling efficiency of AT2 cells was 77% (77% ± 5.40%, n=4) (Figure S1b). In this mouse line, 4.5% of Tom^pos^/total cells were labeled in mice exposed to normal water, indicating leakiness. This percentage is increased to 20.5% of Tom^pos^/total cells in mice exposed to tamoxifen water for one week (Figure S2 a-d) indicating robust induction of labelling. Next, the distribution of fluorescence intensity of the tdTomato^Pos^ cells was analyzed on lung sections. The threshold was set at 22% based on the flow cytometry data and then the intensity of Sftpc immunofluorescence staining was quantified in each cell located on the left side (Tom^Low^) and the right side (Tom^High^) of the threshold. Both populations contain Sftpc^Low^ and Sftpc^High^ cells, thereby suggesting heterogeneity of both Tom^Low^ and Tom^high^ subpopulations in terms of Sftpc level (Figure 1b). Interestingly, PCR from the genomic DNA for the presence of the STOP codon in the *Rosa26* locus (which is normally removed upon induction of Cre activity) indicated that the LoxP-STOP-LoxP site was partially recombined in Tom^Low^ cells versus Tom^High^ cells (Figure 1c). This incomplete deletion could explain the presence of these two subpopulations of tdTomato cells. Next, we investigated whether incomplete deletion of the STOP codon in the [*Sftpc^CreERT2/+^; tdTomato^flox/flox^*], allowing in the Tom^Low^ population only one Tomato copy to be expressed against two copies for the Tom^High^ cells, was the sole reason for the difference in the expression of *Tomato*. To test this possibility, we used [*Sftpc^CreERT2/+^; tdTomato^flox/+^*] mice, with only one copy of Tomato. In case of insufficient recombination, only one population of Tomato positive cells should be observed. Upon treatment with tamoxifen, our flow cytometry results indicate the presence of Tom^Low^ and Tom^High^ subpopulations in these mice as well (Figure S1 c-d) supporting our conclusion that the existence of two subpopulations in [*Sftpc^CreERT2/+^; tdTomato^flox/flox^*] lungs, based on different levels of *Tomato* expression, results from the differential expression of Tomato from the *Rosa26* promoter as well as from the incomplete recombination of the STOP codon in the *Rosa26* allele in Tom^High^ vs. Tom^Low^ cells.

**Figure 1:**
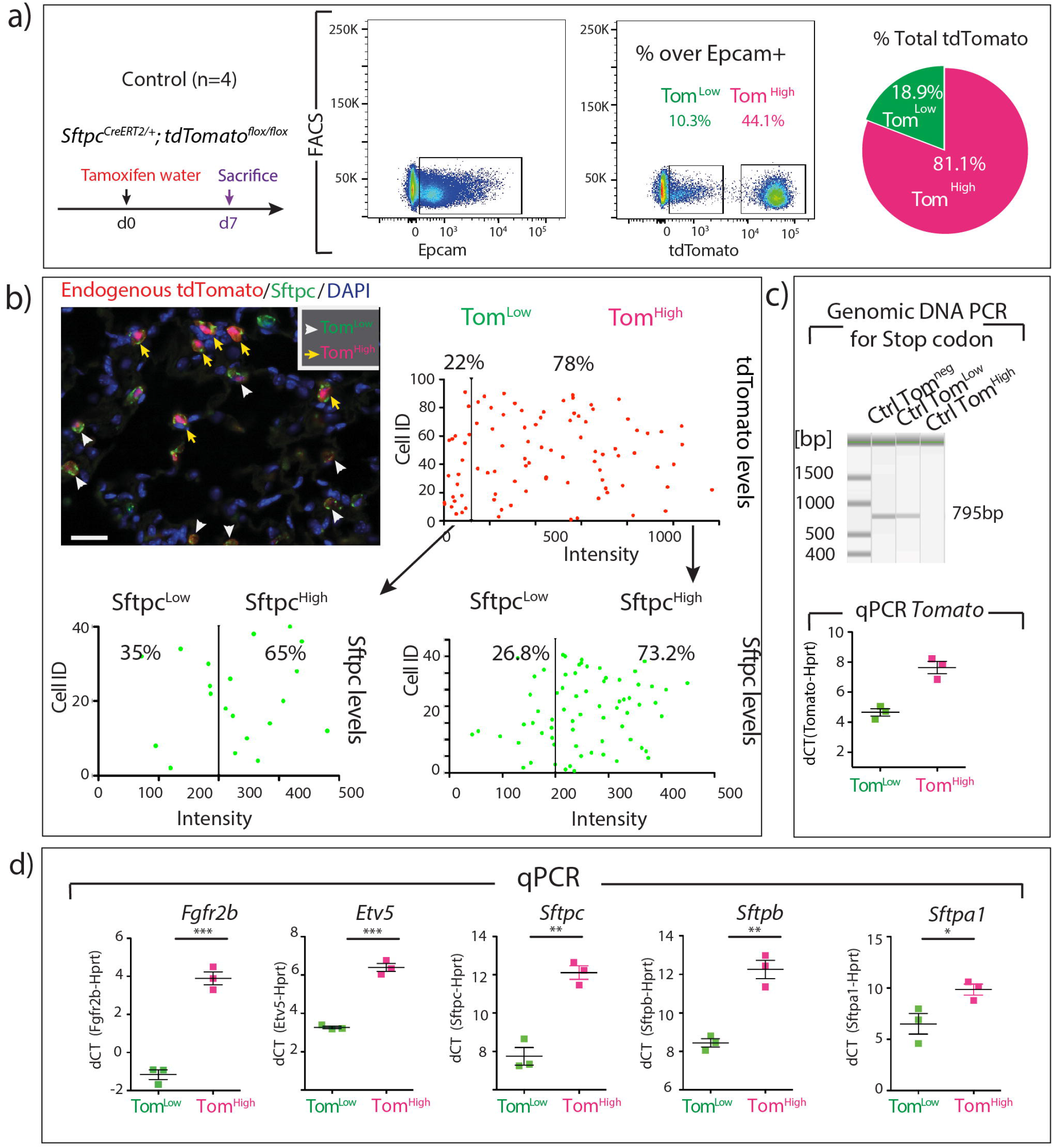
Identification of two populations of AT2-lineage labeled cells, named tdTomato^Low^ (Tom^Low^) and tdTomato^High^ (Tom^High^). **a)** Timeline of tamoxifen treatment of Sftpc^CreERT2/+^, *tdTomato^flox/flox^* mice (n=4) and representative flow cytometry of EpCAM positive population selection and identification of Tom^Low^ (10.3%) and Tom^High^ (44.1%) populations based on the tdTomato level. Pie chart shows the percentage of Tom^Low^ (18.9%) and Tom^High^ (81.1%) in total tdTomato positive cells. **b)** Representative Sftpc immunofluorescence staining and localization of Tom^Low^ and Tom^High^ in the alveoli (Scale bar: 50μm). Plots display quantification of tdTomato fluorescent intensity of tdTomato positive cells and further quantification of Sftpc level in Tom^Low^ and Tom^High^ populations separately. **c)** PCR on genomic DNA isolated from FACS-based sorted Tom^Low^ and Tom^High^ cells and qPCR analysis of FACS-based sorted Tom^Low^ and Tom^High^ cells for the expression of *Tomato*. **d)** qPCR analysis of FACS-based sorted Tom^Low^ and Tom^High^ cells for the expression of *Fgfr2b*, *Etv5*, *Sftpc*, *Sftpb* and *Sftpa1*. Data are presented as mean values ± SEM. *p < 0.05, **p < 0.01, ***p < 0.001.

Next, qPCR was performed on FACS-isolated Tom^Low^ and Tom^High^ cells. Our results show that *Fgfr2b* and its associated downstream target *Etv5*, as well as the differentiation markers *Sftpc, Sftpb*, and *Sftpa1,* were significantly enriched in Tom^High^ vs. Tom^Low^ cells (Figure 1d). Thus, we conclude that Tom^High^ represents the *bona fide* mature AT2 cells and we hypothesized that the lineage-related Tom^Low^ cells represent immature AT2 cells.

### ATAC-seq analysis indicates that Tom^Low^ cells and Tom^High^ cells are distinct cell populations

To analyse the genome-wide profiling of the epigenomic landscape, an assay for transposase-accessible chromatin using sequencing (ATAC-seq) was performed on Tom^Low^ and Tom^High^ subpopulations (Figure 2a). Common and distinct peaks were identified for Tom^Low^ and Tom^High^ cells (Figure 2b).

**Figure 2:**
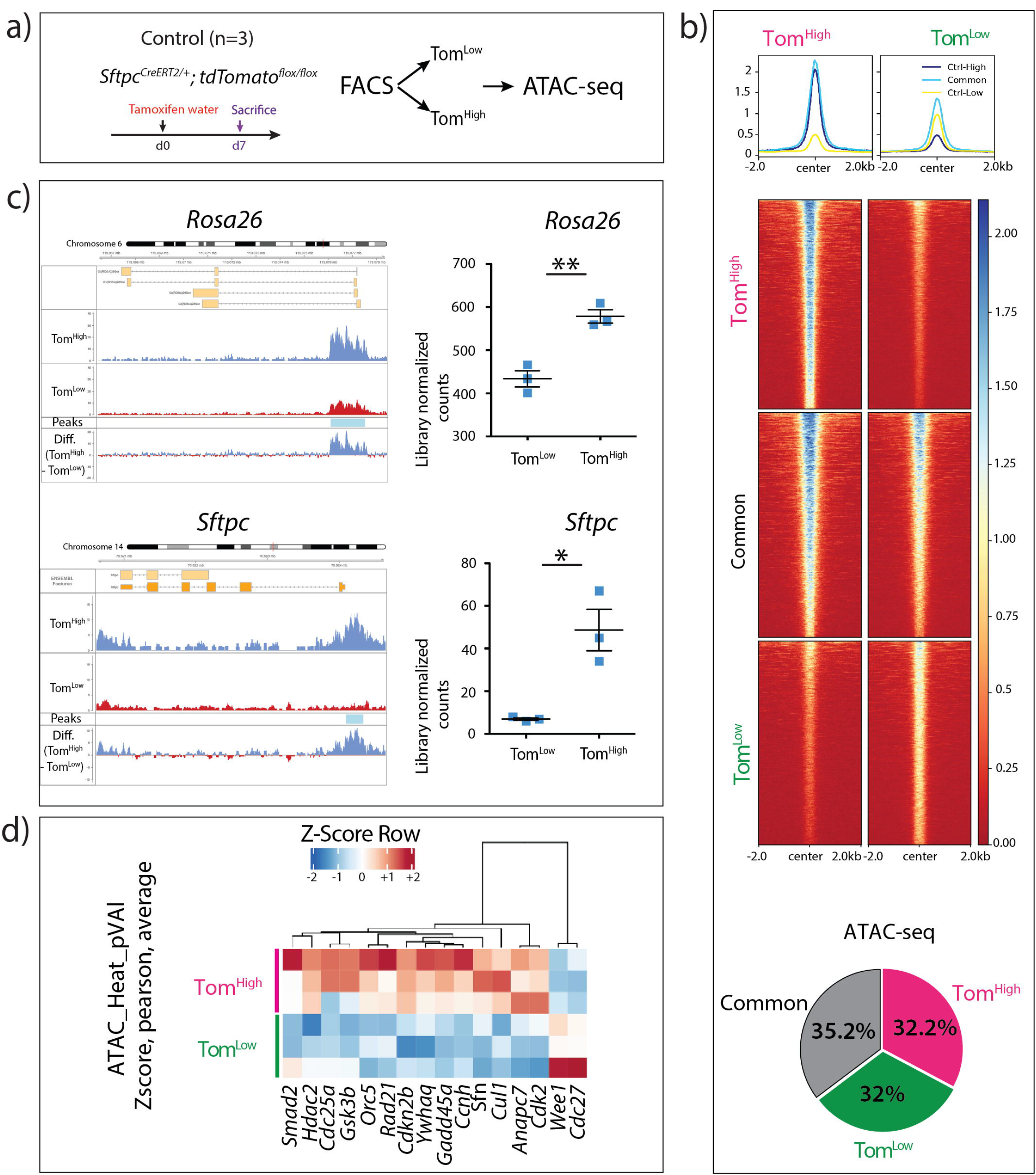
ATAC-seq analysis on FACS-based sorted Tom^Low^ and Tom^High^ populations. **a)** Timeline of tamoxifen treatment of Sftpc^CreERT2/^, *tdTomato^flox/flox^* mice (n=3). ATAC-seq was carried on FACS-based sorted Tom^Low^ and Tom^High^ populations. **b)** Coverage heat maps of Tom^Low^ and Tom^High^, displaying genome-wide regions of differential open chromatin peaks in Tom^Low^ versus Tom^High^. Tom^Low^ chromatin is less open and transcriptionally less active compared to Tom^High^. ATAC-seq analysis of peaks based on the cutoffs shows 3605 up-regulated in Tom^High^ (FDR < 0.05, log2FC > 0.585, baseMean > 20), 3512 up-regulated in Tom^Low^ (FDR < 0.05, log2(FC) > 0.585, baseMean > 20) and 3878 non-regulated (baseMean > 20, FDR > 0.5, log2(FC) between −0.15 and 0.15) which means 32% and 32.8% of the genome is differently accessible in Tom^Low^ and Tom^High^, respectively. **c)** ATAC-seq histogram of average read coverage at *Rosa26* locus shows distinct ATAC-seq peaks at the promoter and denser chromatin in Tom^Low^ compared to Tom^High^ in this locus. Reprehensive peaks of Tom^Low^ and Tom^High^ are the average of three independent samples and the graph shows the quantification of peaks at *Rosa26* locus. **c)** ATAC-seq profile at *Sftpc* locus shows distinct ATAC-seq peaks at the promoter and denser chromatin in Tom^Low^ compared to Tom^High^. Reprehensive peaks of Tom^Low^ and Tom^High^ are averages of three independent samples and the graph displays the quantification of peak regions of Sftpc locus. The ATAC-seq data have been normalized for sequencing depth and the scale on the y-axis was chosen for optimal visualization of peaks **d)** Heatmap based on the ATAC seq data of top cell cycle-related genes differentially regulated in Tom^Low^ and Tom^High^. FDR: the significance of results by Benjamini-Hochberg correction of multiple tests (n=3).

Further analysis of the ATAC-seq data using the Reactome database indicated that chromatin in loci of genes belonging to metabolism, cholesterol metabolism, surfactant mestabolism and triglyceride biosynthesis was more open in Tom^High^ cells. These data agree with the known role of mature AT2 cells in surfactant production. On the other hand, Tom^Low^ cells exhibit more accessibility for genes belonging to immune system, both adaptive and innate as well as extracellular matrix (ECM) organization, ECM proteoglycans and degradation of the ECM. These results suggest a new and important function for the Tom^Low^ in interacting with the immune system, potentially displaying also migratory capabilities through ECM degradation (Figure S3).

Our ATAC-seq data also agree with the decrease in *Tomato* (expressed from the *Rosa26* promoter) and *Sftpc* expression at the mRNA level in Tom^Low^ vs. Tom^High^. In the *Rosa26* and *Sftpc* loci (Figure 2c), the peaks corresponding to the open chromatin configuration were detected at much higher level in Tom^High^ vs. Tom^Low^ and this difference was confirmed to be statistically significant upon quantification (Figure 2c). Interestingly, the analysis of ATAC-seq data also suggests that Tom^Low^ cells display reduced expression of cell cycle genes compared to Tom^High^ cells (Figure 2d). This decrease in cell cycle genes was confirmed by the analysis of our gene array data between Tom^Low^ and Tom^High^ (data not shown). Overall our data indicate that in homeostatic conditions the Tom^Low^ cells fit the profile of a quiescent population.

### AT2-Tom^Low^ and Tom^High^ display different colony-forming capabilities

To compare the self-renewal capacity of Tom^Low^ and Tom^High^, FACS-based sorted cells were co-cultured with CD31^neg^CD45^neg^Epcam^neg^Sca1^pos^ resident lung mesenchymal cells according to a previously published protocol (Figure 3a). Tom^High^ behaved as *bona fide* AT2 cells as they formed alveolospheres with the expected colony-forming efficiency (Figure 3b-d), whereas Tom^Low^ displayed very weak organoid forming capabilities, which is in line with their proposed quiescent status. Both populations transdifferentiated into RAGE-positive AT1 cells after 14 days in culture (Figure 3c).

**Figure 3:**
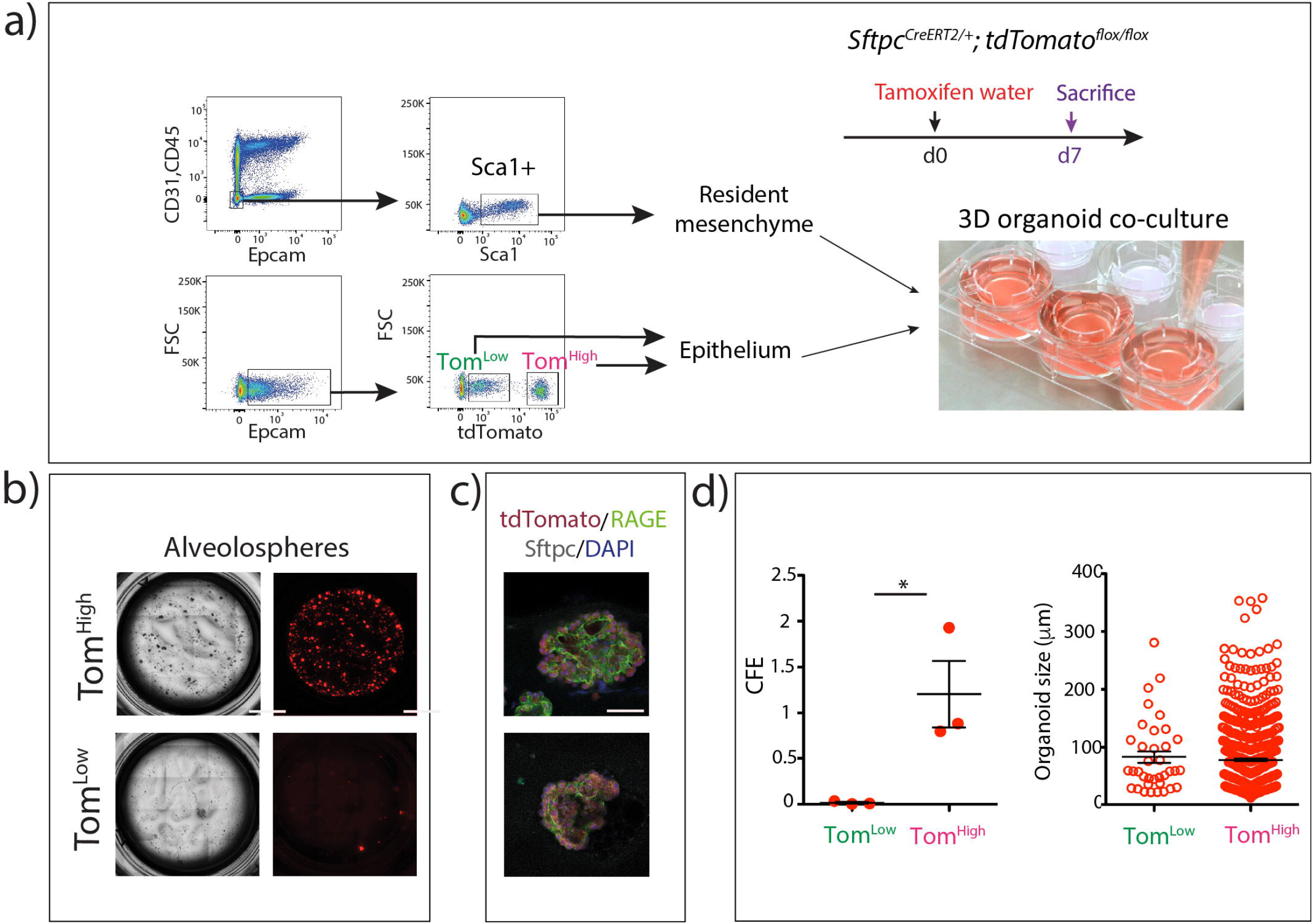
Different colony-forming capabilities of Tom^Low^ and Tom^High^. **a)** Representative flow cytometry shows the gating strategy of CD31^Neg^CD45^Neg^Epcam^Neg^population and further selection of Sca1+ resident mesenchymal cells, as well as the selection of Tom^Low^ and Tom^High^ from Epcam^pos^ population. Mesenchymal cells were co-cultured with Tom^Low^ and Tom^High^ separately (n=3). **b)** Representative alveolospheres from Tom^Low^ and Tom^High^ (n=3) (Scale bar: 100μm). **c)** Representative Sftpc and Rage Immunofluorescent staining of alveolospheres after 14 days in culture (Scale bar: 50μm). **d)** Quantification of Colonyforming efficiency (CFE) and alveolospheres size in Tom^High^ and Tom^Low^ (n=3).

### Expansion of Tom^Low^ population following pneumonectomy

A critical question regarding these two populations of AT2 cells is whether they are differentially engaged in the context of lung regeneration. We, therefore, used the mouse pneumonectomy model to trigger lung regeneration, a process that is tightly associated with the proliferation of AT2 cells [18][19]. [*Sftpc^CreERT2/+^; tdTomato^flox/flox^*] mice (n= 4) were treated with tamoxifen water for one week to label the AT2 lineage. The mice were then put for 2 weeks on normal water to ensure enough time for tamoxifen clearance. Unilateral left lung pneumonectomy (PNX) was then performed to induce the process of compensatory growth in the remaining right lobes. Control mice (Sham) underwent the same process but without removal of the left lobe (Figure 4a). The animals were euthanized at day 7 post-surgery and the lungs were processed for FACS. The quantification of the abundance of the Tom^Low^ and Tom^High^ over the total number of Epcam^Pos^ cells in Sham and PNX indicates that the ratio of Tom^High^ cells/total number of Epcam^Pos^ cells remained unchanged between the two conditions, while the ratio of Tom^Low^ cells was significantly increased in the context of PNX versus Sham (Figure 4b). This suggest that Tom^Low^ cells rather than the previously thought Tom^High^ cells are the ones contributing to the process of lung regeneration. These two AT2 populations were isolated by FACS for further analysis by qPCR. Upon PNX, the expression of *Fgfr2b*, *Etv5*, *Sftpc*, as well as *Cyclin D1* (*Ccnd1*) and *Cyclin D2* (*Ccnd2*) was significantly upregulated in Tom^Low^. A trend towards an increase was also observed in Ki67 expression (Figure 4c). These results are consistent with Fgfr2b signaling activation and proliferation in Tom^Low^ cells in the context of lung regeneration. By contrast, Tom^High^ cells showed no difference in *Fgfr2b*, *Etv5* and *Sftpc* between PNX and Sham conditions. Surprisingly, we noticed an increase in *Ki67*, *Ccnd1* and *Ccnd2* (Figure 4d) but as the number of Tom^High^ cells is not increased in PNX vs. Sham, the significance of these results is not clear.

**Figure 4:**
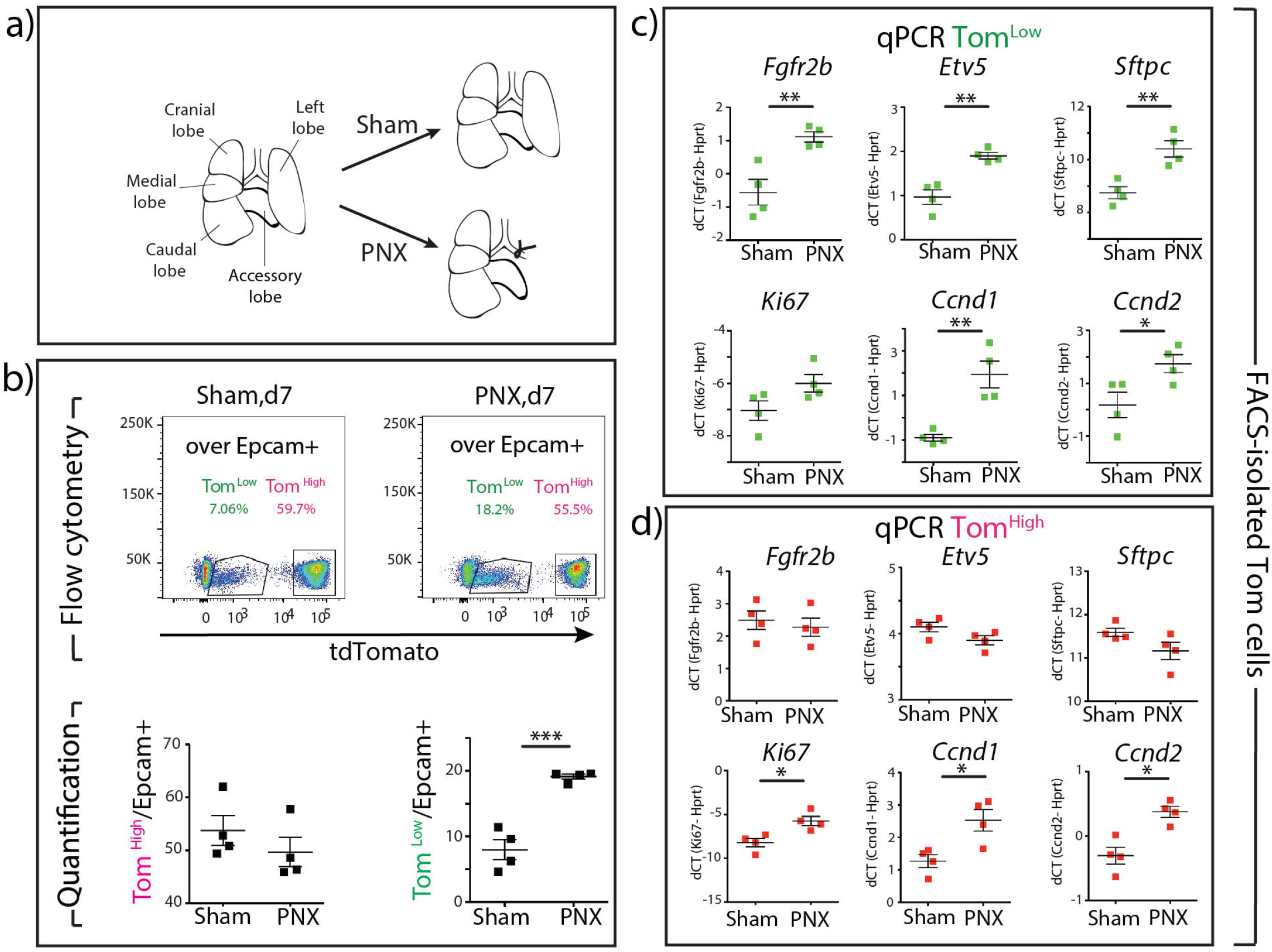
Expansion of Tom^Low^ but not Tom^High^ following pneumonectomy. **a)** Schematic representation of PNX and sham models. **b)** Representative flow cytometry analysis of Tom^Low^ and Tom^High^ populations 7 days after PNX and sham and the quantification of Tom^Low^ and Tom^High^ percentages in Epcam^Pos^ population between sham and PNX groups (n=4) (Scale bar: 250μm). **c)** qPCR analysis of FACS-based sorted Tom^Low^ population for *Fgfr2b*, *Etv5*, *Sftpc, ki67, Ccnd1* and *Ccnd2* expressions. **d)** qPCR analysis of FACS-based sorted Tom^High^ population for *Fgfr2b*, *Etv5*, *Sftpc, ki67, Ccnd1* and *Ccnd2* expressions (Scale bar: 250μm and 50μm). Data are presented as mean values ± SEM. * p < 0.05, ** p < 0.01, *** p < 0.001.

### Loss of Tom^High^ cells leads to expansion of Tom^Low^ cells

We made use of the precision-cut lung slices (PCLS) to follow the fate of tdTomato^Pos^ cells over time *in vitro*. Our flow cytometry results indicate that this approach leads to the drastic loss of the Tom^High^ cells (Figure 5a). Therefore, we took advantage of this system to monitor the fate of Tom^Low^ cells overtime after a massive loss of Tom^High^. Over a culture period of 4 days, we observed the formation of cell clusters and a significant increase in tdTomato intensity (Figure 5b,c), suggesting the differentiation of the Tom^Low^ cells towards mature (Tom^High^) AT2 cells. We also noted the presence of clusters of tdTomato^Pos^ cells at day 10 (Figure 5d). In the future, innovative strategies to carry out the lineage tracing of Tom^Low^ cells will have to be developed to characterize their contribution to the Tom^High^ pool. Therefore, our results agree with our previous observation that Tom^Low^ cells are capable to proliferate in the context of PNX.

**Figure 5:**
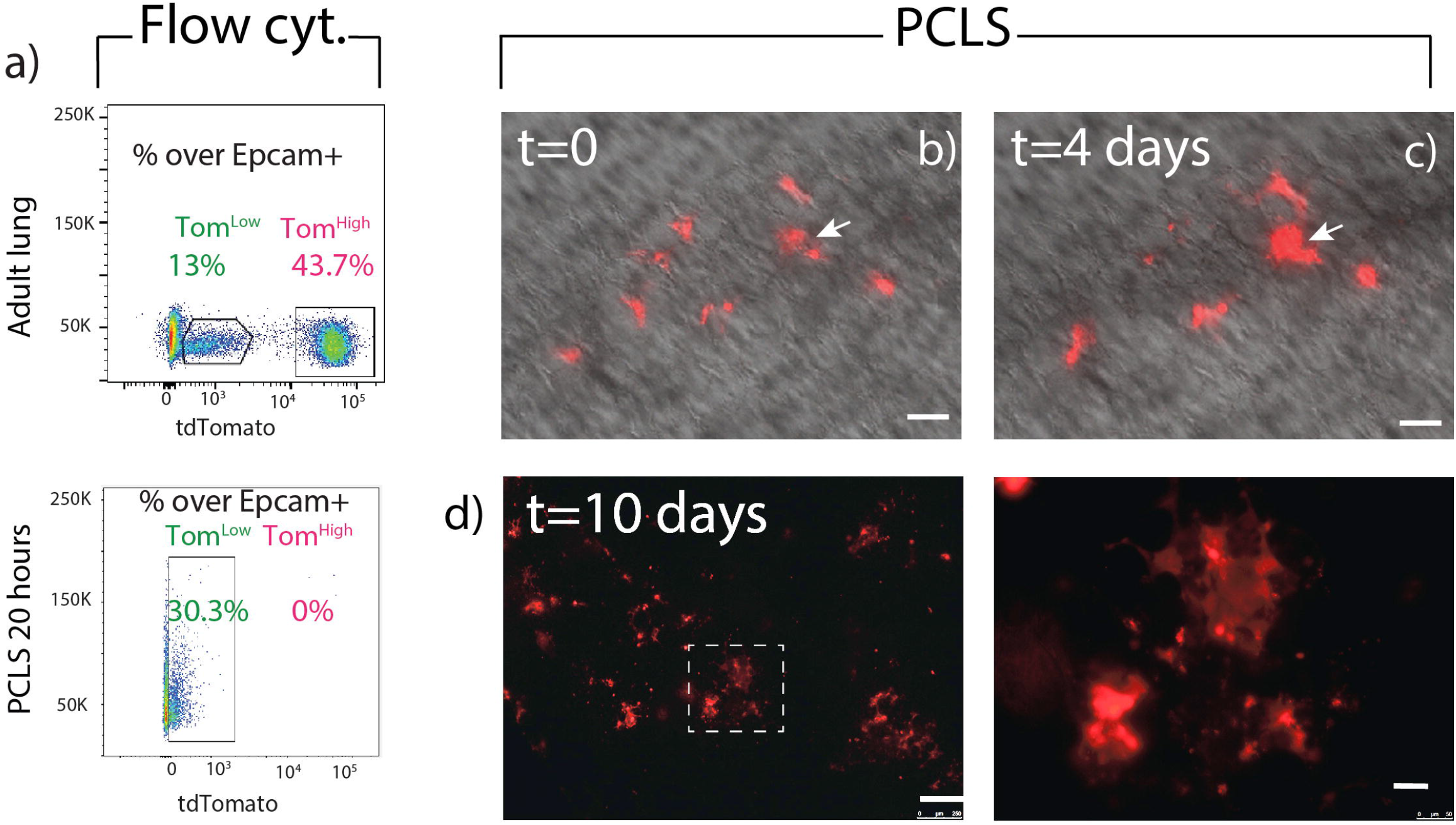
Characterization of the fate of Tom^Low^ cells in precision-cut lung slices (PCLS). **a)** Representative flow cytometry analysis of Tom^Low^ and Tom^High^ before processing the lungs for PCLS and in freshly generated PCLS. **b)** Visualization of the Tom^Low^ in PCLS at t=0. **c)** Visualization of the Tom^Low^ in PCLS at t=4 days. **d)** Visualization of the Tom^Low^ in PCLS at t=10 days. Low and high magnification. Scale bar 250 μm (low magnification) and 50 μm (high magnification).

### PD-L1 is a specific surface marker enriched in Tom^Low^

To identify markers differentially expressed between Tom^Low^ and Tom^High^, a gene array was performed on FACS-isolated Tom^Low^ and Tom^High^ cells (data not shown). Among the 100 top genes differentially expressed in Tom^Low^ vs. Tom^High^ cells three cell surface markers namely, *Cd33*, *Cd300lf*, and *PD-L1 (Cd274), all of* which were enriched in Tom^Low^ compared to Tom^High^, were identified. qPCR analysis confirmed the significantly higher expression of *Cd33* and *PD-L1* expressions in Tom^Low^ compared to Tom^High^ (Figure 6a). PD-L1 seems to be an interesting marker as it is an immune inhibitory receptor ligand and its expression is highly increased in adenocarcinoma [14][20]. The use of this marker is also relevant as our ATAC-seq analysis revealed that Tom^Low^ cells are enriched in genes belonging to the immune system (see Fig. S3). The use of *PD-L1*, as a surface marker enriched in Tom^Low^ cells, was further validated using PD-L1 immunofluorescence staining and flow cytometry. To this end, PD-L1 immunofluorescence staining on cytospins confirms a higher level of protein at the plasma membrane in Tom^Low^ compared to Tom^High^ (Figure 6b). Moreover, flow cytometry analysis of Tom^Low^ and Tom^High^ populations separately showed that 46.9% of Tom^Low^ cells were PD-L1^Pos^ while only 0.77% of Tom^High^ cells were PD-L1^Pos^ (Figure 6c). In addition, PD-L1 immunofluorescence staining on lung sections localize tdTomato^Pos^ PD-L1^Pos^ cells in the alveoli (Figure 6d). In conclusion, these results suggest that PD-L1 antibodies could be instrumental in differentially isolating the equivalent of Tom^Low^ vs. Tom^High^ in wild type lungs.

**Figure 6:**
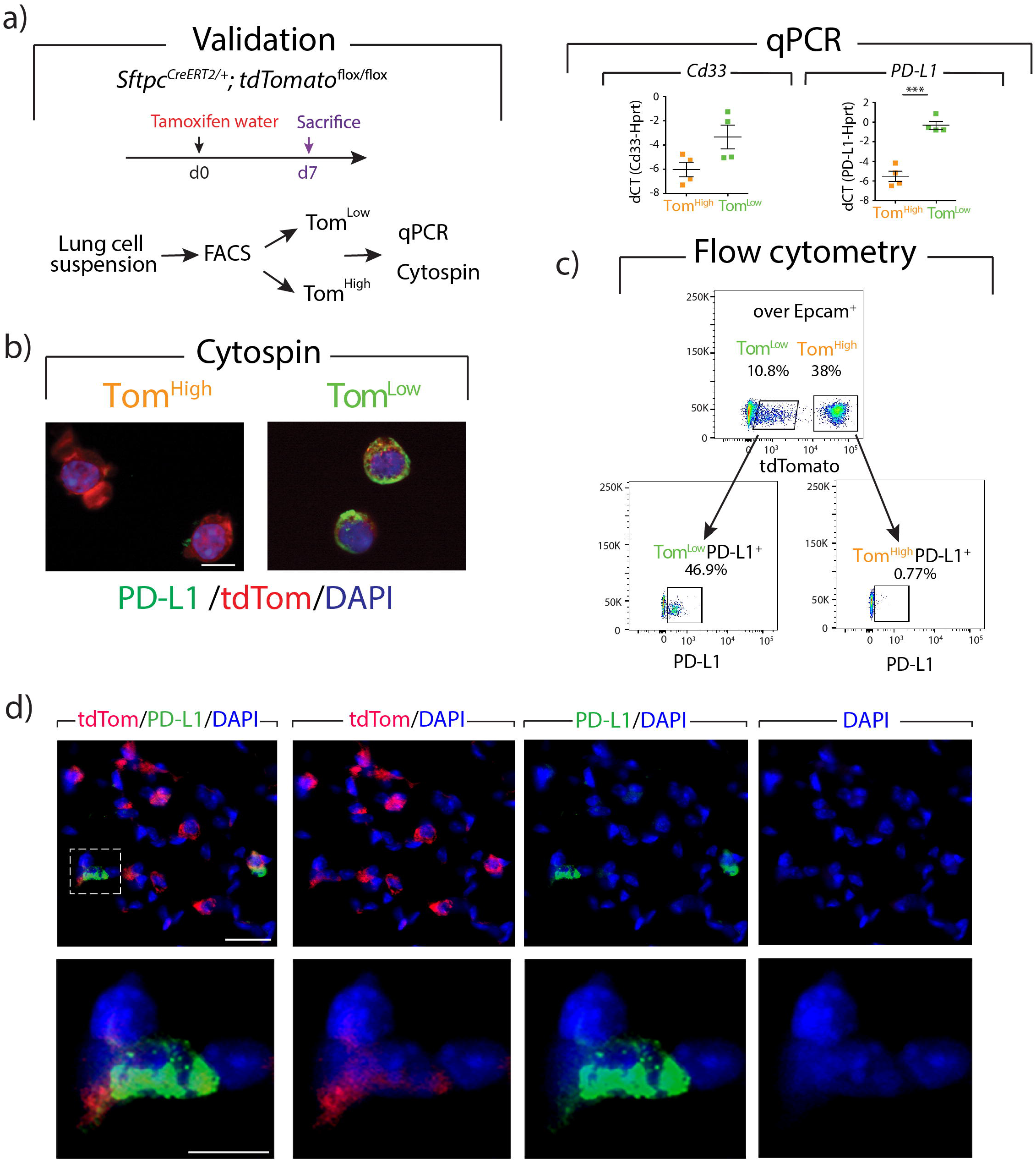
PD-L1 is a specific surface marker enriched in Tom^Low^. **a)** Validation of acquired gene array data with qPCR for the expression level of *Cd33* and *PD-L1* in Tom^Low^ compared to Tom^High^. Data are presented as mean values ± SEM. *p < 0.05, **p < 0.01, ***p < 0.001. **b)** Representative PD-L1 Immunofluorescent staining on Tom^Low^ and Tom^High^ cytospin cells (Scale bar: 50μm). **c)** Representative flow cytometry analysis of PD-L1^Pos^ population in Tom^Low^ and Tom^High^. d) Representative PD-L1 Immunofluorescent staining on the lung sections (scale bar: 10μm)

### Identification of AT2 PD-L1^+^ Sftpc^Low^ in the wild type mice

To address whether PD-L1 could be used to isolate the equivalent of Tom^Low^ and Tom^High^ without the need for a lineage tracing approach, FACS-based analysis was performed on isolated C57BL/6 lungs. AT2 cells selection was done based on the gating of Epcam^Pos^, Cd49f^Inter.^, Podoplanin^Neg^ population, from which the percentage of PD-L1 positive and negative cells were analyzed (Figure 7a). In average, out of the AT2 population, 8.99% ± 0.51% (n=4) of AT2 cells are PD-L1^Pos^ and 90.23% ± 0.58% (n=4) are PD-L1^Neg^ (Figure 7a). This ratio is consistent with the finding that most (80%) of the lineage-traced AT2 cells were composed of Tom^High^ (PD-L1^Neg^) cells (Figure 1a). qPCR analysis on sorted PD-L1^Pos^ and PD-L1^Neg^ AT2 cells indicates a higher level of *Fgfr2b, Etv5*, and *Sftpc* in PD-L1^Neg^ compared to PD-L1^Pos^ cells (Figure 7b). This result is also in line with PD-L1^Neg^ cells correlating with Tom^High^ cells, while the expression profile of PD-L1^High^ fits with these cells being the equivalent of the Tom^Low^. Finally, flow cytometry analysis of alveolar epithelial cells (Epcam^Pos^, Cd49f^inter^) stained with Sftpc and PD-L1 identified a subpopulation of AT2 cells (6.8%) displaying low level of Sftpc and high level of PD-L1 (Figure 7c). This Sftpc^Low^ PD-L1^High^ likely contains the Tom^Low^ cells.

**Figure 7:**
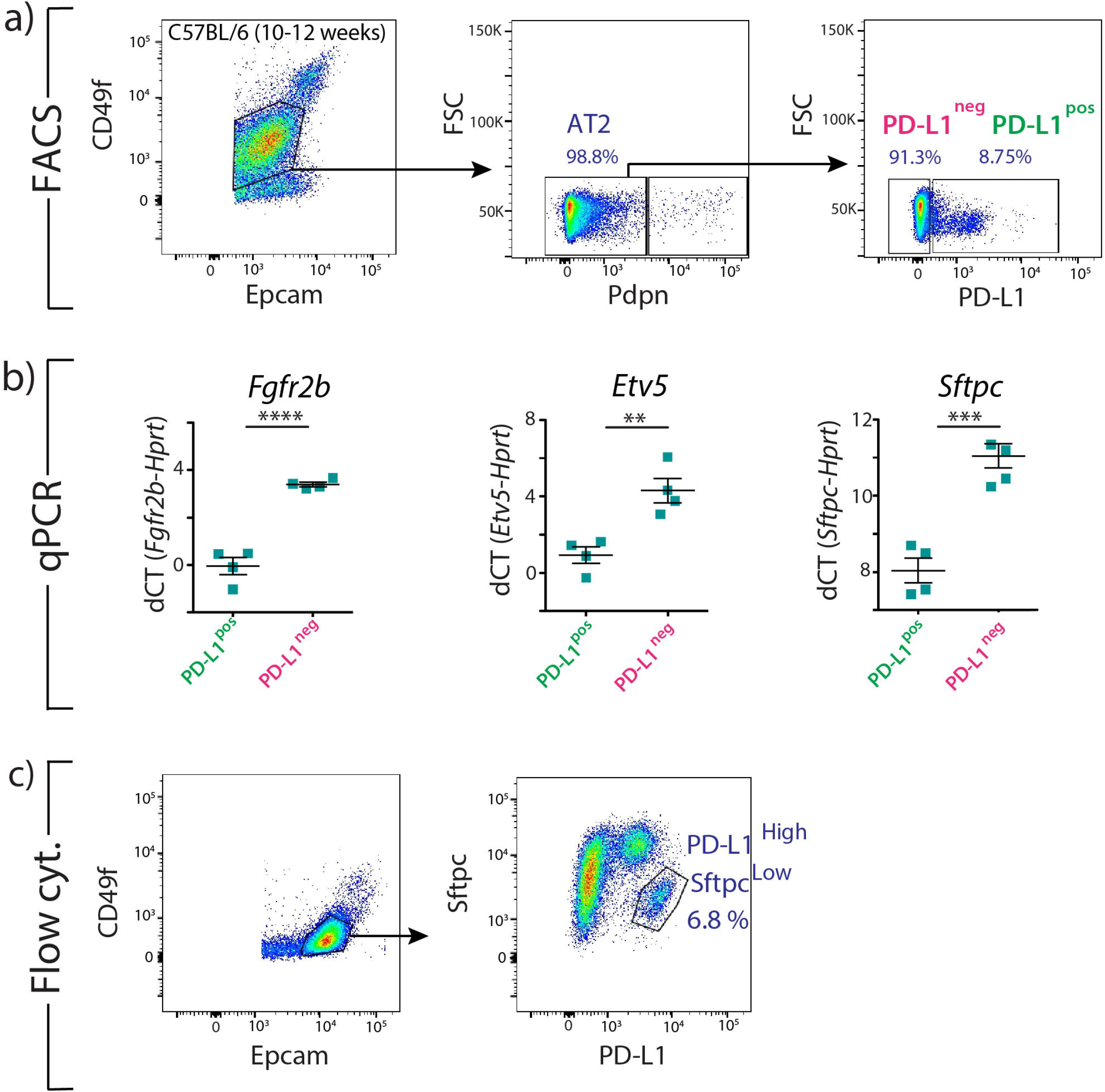
Identification of AT2 PD-L1^+^ Sftpc^Low^ in the wild type mice. **a)** Representative flow cytometry analysis of wild type mice lungs showing the gating strategy of Epcam^+^ CD49f^Inter^ population, followed by negative selection of AT2 cells with the exclusion of AT1 Pdpn^+^ cells. Representative flow cytometry analysis of AT2 cells based on PD-L1 marker. **b)** qPCR analysis of FACS-based sorted PD-L1^Pos^ and PD-L1^Neg^ for *Fgfr2b*, *Etv5*, and *Sftpc* expressions (n=4). Data are presented as mean values ± SEM. *p < 0.05, **p < 0.01, ***p < 0.001. **c)** Representative flow cytometry analysis of Sftpc intracellular staining shows a distinct Sftpc^Low^ PD-L1^High^ population of AT2 cells.

### Decrease of *bona fide* EPCAM^Pos^HTll-280^High^ alveolar type 2 cells and amplification of EPCAM^Pos^HTll-280^Low/Neg^ PD-L1^High^ cells in human end-stage idiopathic lung fibrosis compared to donor lungs

Idiopathic pulmonary fibrosis (IPF) is a disease related to AT2 cell deficiency [21] [22][23]. We used flow cytometry to analyze the epithelial component of donor (n=5) and IPF (n=5) lungs (Figure 8a). From the live cells, we first excluded the CD31^Pos^CD45^Pos^ and then selected the EPCAM^Pos^ cells for further analysis using the human AT2 cell marker HTll-280 and the surface marker PD-L1. Our data indicate that HTll-280^High^ PD-L1^Neg^ (Q1), representing the *bona fide* differentiated AT2 cells, were drastically reduced in the context of IPF (25.21% ± 12.46 vs. 80.66% ± 3.49). More interestingly, the number of HTll-280^Low/Neg^ PD-L1^High^ (Q3) was drastically increased (13.42% ± 4.92 vs. 1.13% ± 0.47) (Figure 8b), suggesting that HTll-280^Low^ PD-L1^High^ epithelial cells could represent a pool of progenitors linked to the deficient AT2 lineage. The comparison of the expression of *SFTPC* and *SCGB1A1* for cells in Q1 (AT2 cells), Q3 (potential equivalent to Tom^Low^ cells) and Q4 (non-AT2 cells, mostly bronchial epithelial cells) in Donor and IPF lungs was carried out by qPCR (Figure 8c). Regardless of the IPF or Donor origin of these cells, our results indicated that Q3 (potential equivalent to Tom^Low^ cells) express less *SFTPC* than Q1 (AT2 cells) confirming the flow cytometry data. In addition, Q4 (non-AT2 cells) express more *SCGB1A1* than Q3 (potential equivalent to Tom^Low^ cells) suggesting that Q3 cells do not represent bone fide bronchial cells. Interestingly, in IPF, the difference between *SCGB1A1* expression in Q3 (equivalent to Tom^Low^ cells) and Q1 (AT2 cells) is not statistically significant. Based on these results, we propose that these HTll-280^Low^PD-L1^High^ (Q3) epithelial cells could represent the human equivalent of the Tom^Low^ cells characterized in mice (Figure 8d). Indeed, a similar approach carried out in mice with FACS-isolated Tom^Neg^ (mostly bronchial cells), Tom^Low^ and Tom^High^ out of the Epcam+ cells indicated an almost identical expression profile for *Sftpc* and *Scgb1a1* as the one observed in human lungs (Figure 8d).

**Figure 8:**
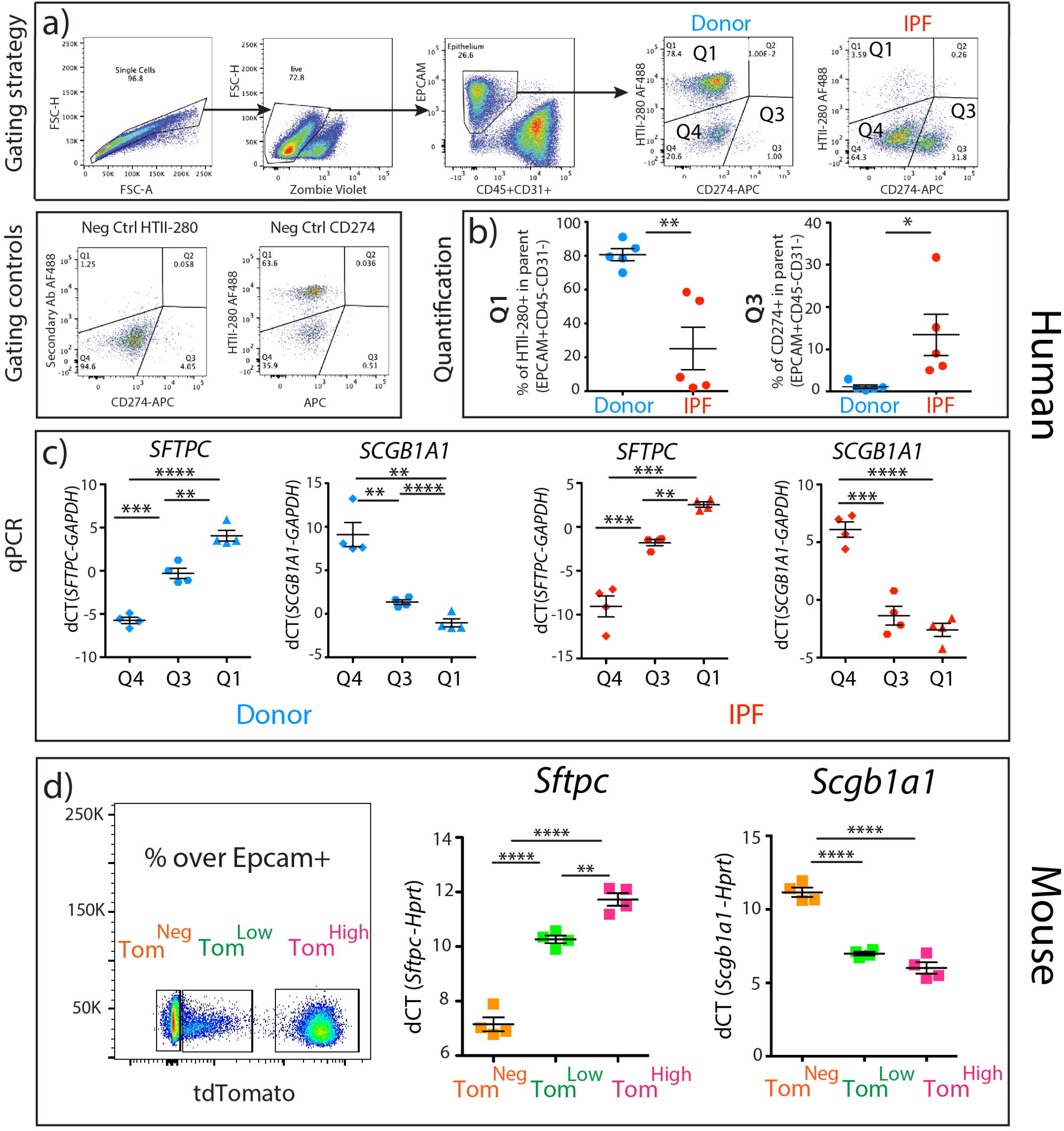
Decrease of bona fide EPCAM^Pos^HTll-280^High^ alveolar type 2 cells and amplification of EPCAM^Pos^HTll-280^Low/Neg^ PD-L1^High^ cells in human end-stage idiopathic lung fibrosis compared to donor lungs. **a)** Representative flow cytometry of single cells and live cells gating, and EPCAM positive population selection and further analysis of HTII-280 and PD-L1 positive populations of IPF and donor lungs. The gating controls panel shows the negative control of HTII-280 and PD-L1 antibodies staining. **b)** Quantification of the number of HTII-280^Pos^ cells PD-L1^Pos^ cells in IPF versus donor (n=5). Data are presented as mean values ± SEM. *p<0.05, **p< 0.01, ***p < 0.001. **c)** qPCR analysis of FACS-based sorted HTll-280^Neg^PD-L1^Neg^ (Q4) HTll-280^Low/Neg^PD-L1^High^(Q3) and HTll-280^Low/Neg^PD-L1^High^(Q1) cells for *SFTPC* and *SCGB1A1* expressions in IPF lungs (n=4). Data are presented as mean values ± SEM. *p < 0.05, **p < 0.01, ***p < 0.001. **d)** Representative flow cytometry analysis of Tom^Neg^,Tom^Low^ and Tom^High^ selection and qPCR analysis of FACS-based sorted Tom^Neg^,Tom^Low^ and Tom^High^ cells for the expression of *Sftpc* and *Scgb1a1*. Data are presented as mean values ± SEM. *p < 0.05, **p < 0.01, ***p < 0.001.

## DISCUSSION

We have identified a subpopulation of AT2 progenitor cells which is distinct from already identified mature AT2 progenitor subpopulations. Tom^Low^ cells are quiescent and immature cells in the steady-state and unlike AT2 Axin2-positive cells express low level of AT2 differentiation markers. Moreover, Tom^Low^ express significantly lower level of *Axin2* compared to Tom^High^ and *Axin2* expression showed no change after pneumonectomy in these cells (Figure S4 a,b). Zacharias et al. showed that AEPs express higher level of lung development and tube development genes than mature AT2. However, based on our gene array data these genes are expressed at lower level in Tom^Low^ versus Tom^High^ (data not shown). This indicates that AEPs cells are part of the Tom^High^ population. The newly identified Tom^Low^ cells are also different from integrin α6β4 population as these cells are negative for *Sftpc* and therefore, the *Sftpc^CreERT2^* driver line cannot lineage trace this population [7][24]. Finally, The Tom^Low^ population most likely does not contain the BASC for several reasons. First, most of the Tom^Low^ are located in the respiratory epithelium and do not display high level of *Scgb1a1* (Figure 8d). Second, single-cell RNA seq data indicate that compared to AT2 cells, BASC expresses a similar level of Sftpc and Fgfr2 [13] indicating that BACSs are likely contained in the Tom^High^ population. Interestingly, lineage tracing of BASC upon injury indicates that these cells are not the sole contributor to newly formed bronchial airway cells after conducting airway injury or AT2/AT1 cells after alveolar injury suggesting that other resident stem cells such as the AT2 Tom^Low^, the AT2 could contribute to the repair process. Furthermore, PD-L1 is highly expressed in Tom^Low^. PD-L1 expression is also increased in adenocarcinoma and appears to be correlated with elevated tumour proliferation and aggressiveness [14][20][25][16][26]. Further investigations are required to elucidate PD-L1 function in Tom^Low^ and their potential interaction with the immune system as well as their contribution to cancer. In the future, it will be important to design dual lineage tracing strategies based on the expression of Sftpc and PD-L1 to specifically label the Tom^Low^ cells.

In conclusion, we identified a novel population of quiescent and immature AT2 progenitor cells with different gene expression profile from mature AT2 cells which are proliferating after PNX and enriched in PD-L1 expression. The equivalent of this population also was identified in human lungs. Further characterization of these cells in homeostatic and repair/regeneration conditions will allow identifying the signaling pathways activated in these cells with the ultimate goal to enhance repair after injury.

## Acknowledgement

We thank the Kerstin Goth for the animal husbandry and genotyping of the animals.

## Grants

S.B. was supported by grants from the Deutsche Forschungsgemeinschaft (DFG; BE4443/1-1, BE4443/4-1, BE4443/6-1, KFO309 P7 and SFB1213-projects A02 and A04), Landes-Offensive zur Entwicklung Wissenschaftlich-Ökonomischer Exzellenz (LOEWE), UKGM, Universities of Giessen and Marburg Lung Center (UGMLC), DZL, and COST (BM1201). DAA acknowledges support from NHLBI (R01HL141856). J.S.Z was funded through a start-up package from Wenzhou Medical University and the National Natural Science Foundation of China (grant number 81472601). S.H. was supported by the UKGM (FOKOOPV), the DZL and University Hospital Giessen and grants from the DFG (KFO309 P2/8; SFB1021 C05, SFB TR84 B9).

## Supplementary Figure captions

**Figure S1:**
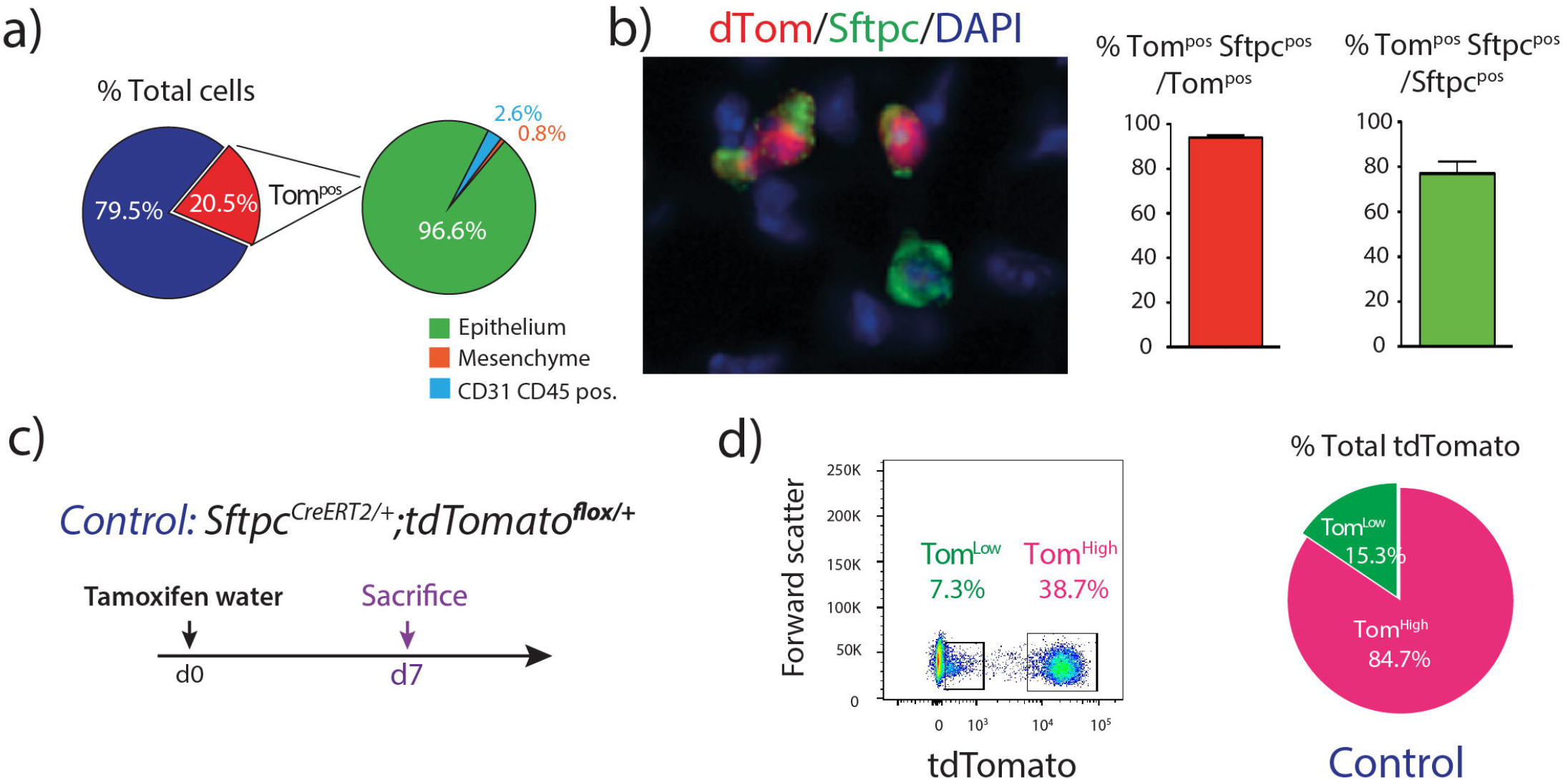
Validation of *Sftpc^CreERT2/+^; tdTomato^flox/flox^* mice. **a)** The pie chart represents the percentage of tdTomato^Pos^ cells in total cells (20.5%) and the percentage of epithelial (96.6%), mesenchymal cells (2.6%) and CD31, CD45 positive cells (0.8%) of the labeled cells. **b)** Representative Sftpc immunofluorescence staining and quantification of tdTomato^Pos^ Sftpc^Pos^ of total tdTomato^Pos^ as well as quantification of tdTomato^Pos^ Sftpc^Pos^ of total Sftpc^Pos^ (n=4). Data are presented as mean values ± SEM. **c)** Timeline of tamoxifen treatment of Sftpc^CreERT2/+^, Tom^flox/+^ mice. **d)** Representative flow cytometry analysis of Tom^Low^ (6.97%) and Tom^High^ (38.7%) of Sftpc^CreERT2/+^, tdTomato^flox/+^ mice. The pie chart shows the percentage of Tom^Low^ (15.3%) and Tom^High^ (84.7%) in total tdTomato positive cells.

**Figure S2:**
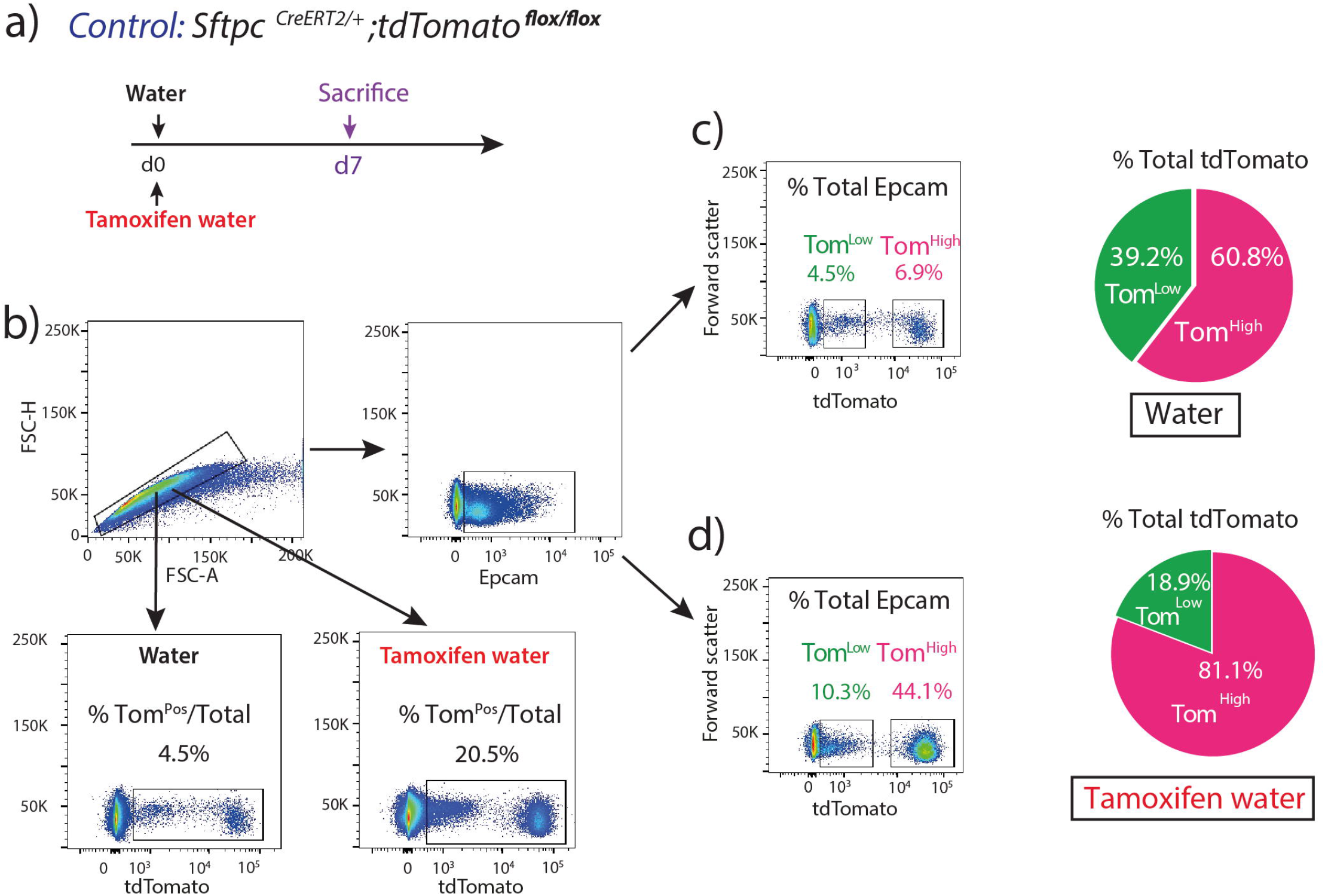
AT2 cells labeling in the absent and the presence of tamoxifen treatment. **a)** Timeline of tamoxifen treatment. One group of Sftpc^CreERT2/+^, *tdTomato^flox/flox^* mice were treated with tamoxifen in drinking water for 7 days and another group no tamoxifen added to the water. **b)** Representative flow cytometry analysis of single-cell selection and further analysis of Epcam^Pos^ population in both groups. **c)** Representative flow cytometry analysis of Tom^Low^ (4.5%) and Tom^High^ (6.9%) in untreated mice with tamoxifen. The pie chart shows the percentage of Tom^Low^ (39.2%) and Tom^High^ (60.8%) in total tdTomato positive cells. **d)** Representative flow cytometry analysis of Tom^Low^ (10.3%) and Tom^High^ (44.1%) in treated mice with tamoxifen. Pie chart shows the percentage of Tom^Low^ (18.9%) and Tom^High^ (81.1%) in total tdTomato positive cells.

**Figure S3:**
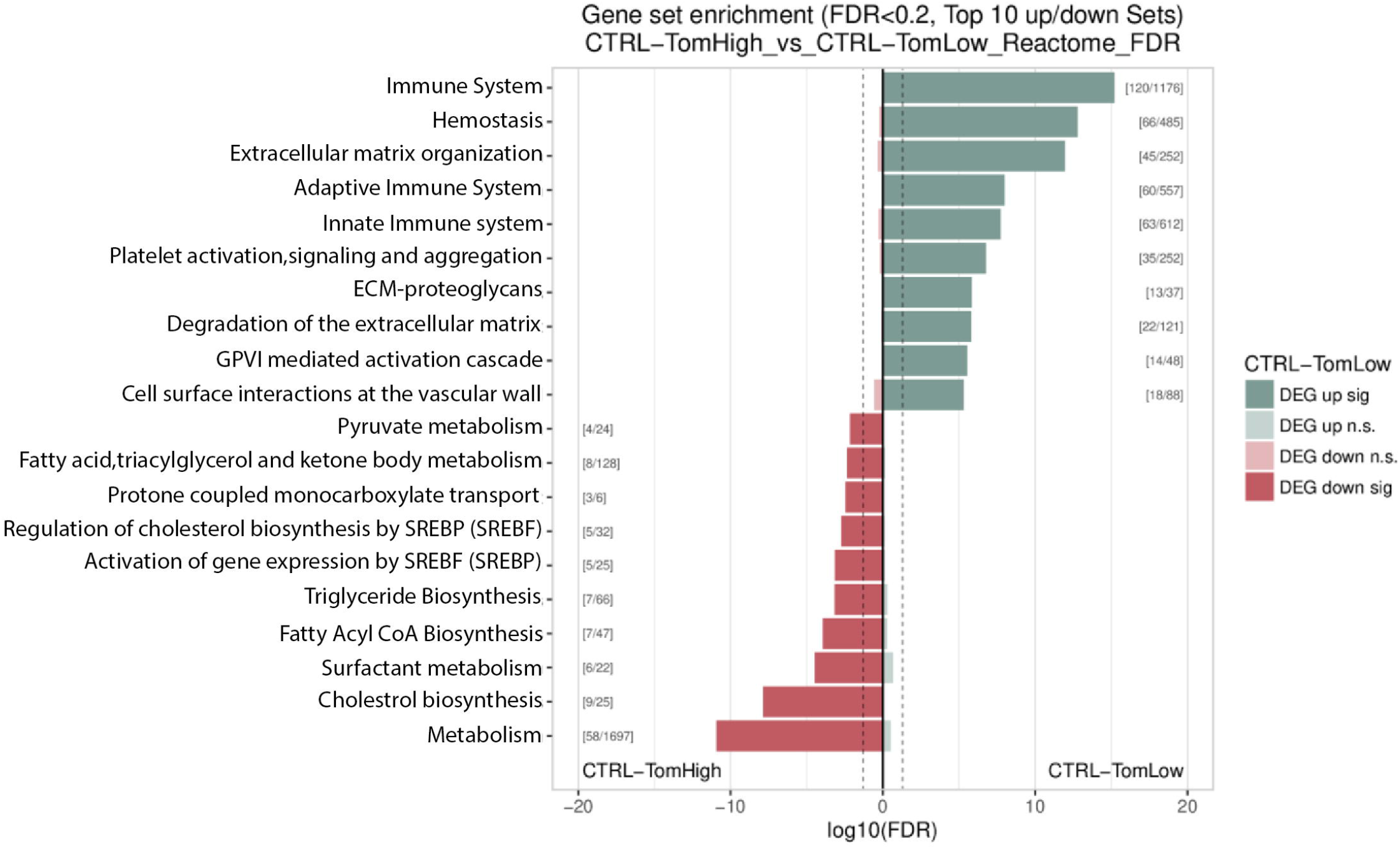
Gene set enrichment for accessible regions in Tom^High^ vs. Tom^Low^. Analysis of peaks obtained in the ATAC-seq experiment for Tom^High^ and Tom^Low^ cells using Kobas for the Reactome database. Peaks overlapping gene body or in close proximity to the transcription starting site of genes were annotated to the corresponding genes. All annotated peaks were split into lists of genes which display more open chromatin in Tom^High^ or Tom^Low^ cells using DESeq2 on unified peak regions. Observed significance was adjusted by Benjamini-Hochberg correction for multiple tests (FDR). The resulting lists were used as input for Kobas to search for enriched terms in different databases. Top 10 terms were chosen by significance (FDR < 0.2). Results indicate that the term immune system is highly enriched in Tom^Low^ cells, indicating that the chromatin of Tom^Low^ cells is more accessible in loci of genes (gene body or promoter) associated with the immune system. Higher accessibility is associated with more transcriptional activity. Numbers in brackets display number of identified genes / total number of genes for term in database. DEG: Differentially expressed genes. Between brackets []: Genes found/total genes in term.

**Figure S4:**
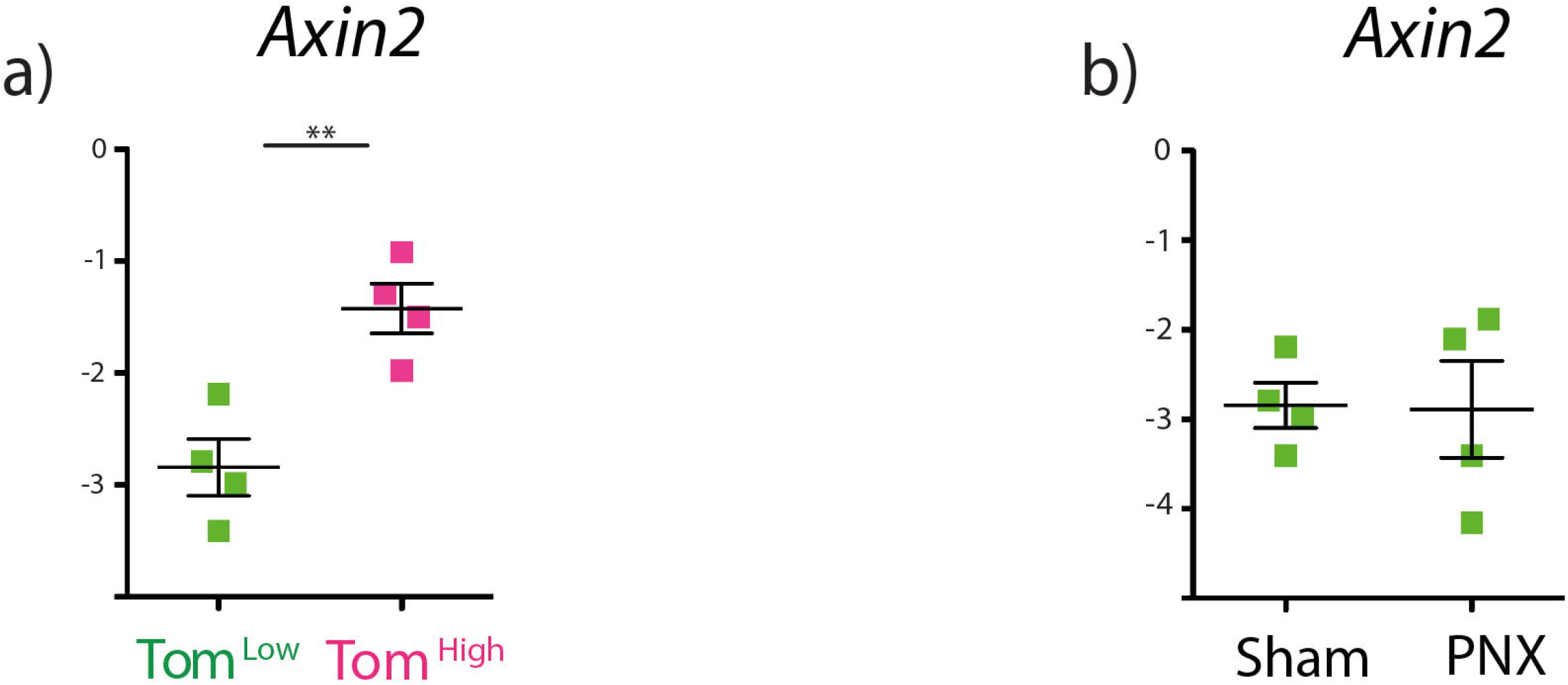
*Axin2* expression analysis. **a)** qPCR analysis of FACS-based sorted Tom^Low^ and Tom^High^ cells for the expression of *Axin2*. **b)** qPCR analysis of FACS-based sorted Tom^Low^ cells for the expression of *Axin2* from sham and PNX groups. Data are presented as mean values ± SEM. *p < 0.05, **p < 0.01, ***p < 0.001.

## Notes

### Competing Interest Statement

The authors have declared no competing interest.

